# Amyloid precursor protein mediates deficits in corticogenesis in Down syndrome cortical organoids

**DOI:** 10.1101/2025.07.26.666926

**Authors:** Karen Rakowiecki, Deepika Patel, Luis Aponte Cofresi, Orly Lazarov

## Abstract

Down syndrome (DS), due to trisomy 21 (T21), occurs in approximately 14.14 per 10,000 live births in the United States. Reduced neural progenitor cell (NPC) proliferation, delayed neurogenesis, impaired cortical lamination and altered cell fate specification are thought to contribute to cognitive impairments in DS individuals. The molecular mechanisms underlying these deficits are not fully understood. Notably, Amyloid precursor protein (APP), located on human chromosome 21 (HSA21), has been extensively implicated in these processes. Mouse models only partially recapitulate DS phenotypes due to genetic, developmental, and species-specific differences. Recent advances in induced pluripotent stem cell (iPSC) derived 3D cortical organoids allow for the study of DS cortical development in a human model system. Here, we show that normalizing APP gene copy number in DS cortical organoids ameliorated deficits in NPC proliferation, neuronal differentiation, and transcriptional programs. Our results demonstrate the value of cortical organoids in uncovering gene-specific roles in DS pathogenesis and identify APP as a promising target for addressing early neurodevelopmental impairments.

## Introduction

DS is one of the most prevalent developmental disorders occurring in 14.14 out of every 10,000 live births in the United States^1^. Studies in the fetal frontal cortex have uncovered reduced NPC proliferation, delayed neurogenesis, and impaired cortical lamination as DS phenotypes^2–6^. Mouse models have recapitulated a number of these deficits^7–11^. Nevertheless, these models are limited in their ability to recapitulate the effect of HSA21 triplication on temporal alterations in neurogenesis^12^. The advent of patient derived iPSCs has allowed for improved elucidation of mechanisms underlying neurodevelopmental impairments in DS. A number of studies in DS NPCs reported deficits in proliferation and cell fate likely stemming from oxidative stress-induced senescence^5,13–22^. How these NPC phenotypes affect downstream cell lineages remains to be fully established. 3D cortical organoids effectively recapitulate aspects of cortical development including generation of NPC layering and diverse neuronal populations^23–26^. Thus, the effect of HSA21 triplication on specific cell populations through corticogenesis can be further elucidated in this model. Limited studies report that DS cortical organoids exhibit reduced expansion, RGC proliferation, and counts of deep and upper layer neurons^27,28^. In one study, transcriptional analysis indicated that layer IV neocortical excitatory neurons are a particularly vulnerable population^28^. Importantly, this model can be utilized to determine how individual HSA21 genes manifest these phenotypes. Knockout of 1 of 3 DSCAM alleles ameliorated a number of deficits^27^. APP is a ubiquitously expressed, single-pass transmembrane protein generated from HSA21^29^. Extensive literature implicates APP in proliferation, cell fate, and migration during cortical development^30–34^. APP gene overdose has been shown to mediate a number of DS CNS phenotypes^35–39^. However, its role in early corticogenesis remains to be investigated. Here, we generated DS cortical organoids to model deficits in early cortical development. We show that DS cortical organoids exhibited deficits in proliferation, neuronal differentiation, and transcription relative to isogenic controls. Notably, silencing of one APP allele mitigated these phenotypes in DS cortical organoids. Taken together, our studies better inform the development of therapeutics targeting cortical development in DS.

## Results

### DS cortical organoids exhibit increased APP protein expression and processing

iPSC lines derived from DS patients and corresponding isogenic controls were obtained from WiCell (Ctrl #1 and DS #1) and National Institute of Neurological Disorders and Stroke (NINDS) Cell and Data Repository (Ctrl #2 and DS #2)^40,41^. To model corticogenesis, iPSC lines were differentiated into dorsal forebrain organoids utilizing a commercially available kit adapted from a previously characterized protocol^42^. Cortical organoids were collected at d25, d50, and d75 to analyze neurogenesis temporally. To determine if HSA21 triplication resulted in increased APP protein expression in cortical organoids, we performed immunoblotting for APP full length (APP-FL). Relative to Ctrl, DS organoids did not exhibit significantly increased APP-FL protein levels at d25 (**Figure 1A-B**). However, DS organoids did exhibit significantly increased APP-FL protein expression at d50 and d75 (**Figure 1C-D**). To determine if APP gene overdose leads to increased APP processing, we performed ELISAs for Aβ in organoid conditioned media. No alterations in Aβ-40 and Aβ-42 levels were observed across genotypes in d25 organoid conditioned media (**Figure 1E-F**). However, d50 and d75 DS cortical organoids exhibited significantly increased secreted Aβ-40 and Aβ-42 relative to Ctrl (**Figure 1E-F**). No alterations in Aβ-42 and Aβ-40 ratio were observed across genotypes (**Figure 1G**).

**Figure 1.**
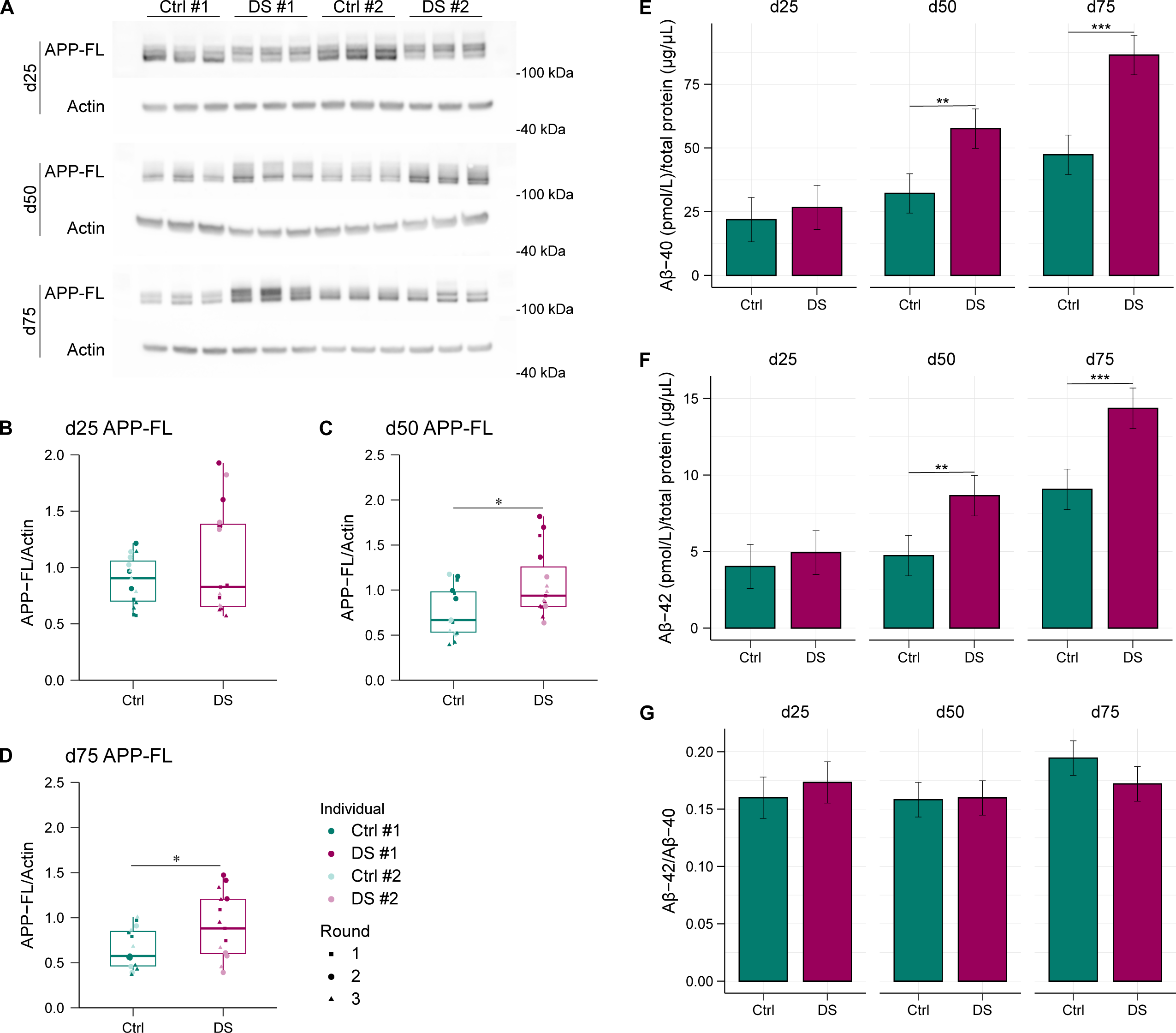
DS cortical organoids exhibit increased expression and processing of APP. (A) Representative immunoblots of APP-FL from d25, d50, and d75 Ctrl and DS dorsal forebrain organoids. (B-D) Quantification of protein expression normalized to β-actin. DS/Ctrl (n = 15, 10 orgs per n, 3 rounds). Data represented as mean ± SEM. Data analyzed by unpaired two-tailed Student’s t-test. * p ≤ 0.05, ** p ≤ 0.01, *** p ≤ 0.001, **** p ≤ 0.0001. (E-G) Levels of secreted Aβ-40 and Aβ-42 and Aβ-42/Aβ-40 ratio from d25, d50, and d75 Ctrl and DS dorsal forebrain organoid conditioned media measured by ELISA. A linear mixed model (LMM) was used to test for differences in Aβ-40, Aβ-42, or Aβ-42/Aβ-40 ratio. Specifically, day (d25, d50, and d75) and line (Ctrl and DS) were fixed effects and individual and round of differentiation were random effects. Post hoc comparisons were performed comparing levels of differences between line within each time point. Multiple comparisons were corrected for using Tukey’s adjustment. DS/Ctrl (n = 9-15, 5-10 orgs per n, 2-3 rounds). Data represented as mean ± SEM. * p ≤ 0.05, ** p ≤ 0.01, *** p ≤ 0.001, **** p ≤ 0.0001.

### DS cortical organoids exhibit reduced expansion and neurogenic protein expression

At d25, proliferative zones model the expanding ventricular zone (VZ) predominantly composed of SRY (sex determining region Y)-box 2 (SOX2)+ radial glia cells (RGCs)^43,44^. To determine the effect of HSA21 triplication on RGC proliferation and organization, immunostaining was performed on Ctrl and DS organoid cryosections for SOX2 and KI67, expressed in higher quantities by proliferating cells^45^. DS proliferative zones exhibited significantly decreased proliferating SOX2+ RGCs relative to Ctrl (**Figure 2A-B**). In continuity, DS proliferative zones were reduced in area and perimeter relative to Ctrl (**Figure 2C-D**). To begin interrogating alterations in neurogenesis, we ran immunoblots for the stemness protein SOX2 and neurogenic proteins doublecortin (DCX) and β-3 tubulin. At d25, DS cortical organoids exhibited significantly increased SOX2 and decreased DCX and β-3 tubulin protein expression relative to Ctrl (**Figure 2E-H**). To determine if deficits in neurogenesis endured, we performed the same immunoblotting at d50. DS cortical organoids exhibited decreased DCX but no alterations in SOX2 and β-3 tubulin protein expression relative to Ctrl suggesting persistent impairments in neuroblasts (**Figure 3A-D**).

**Figure 2.**
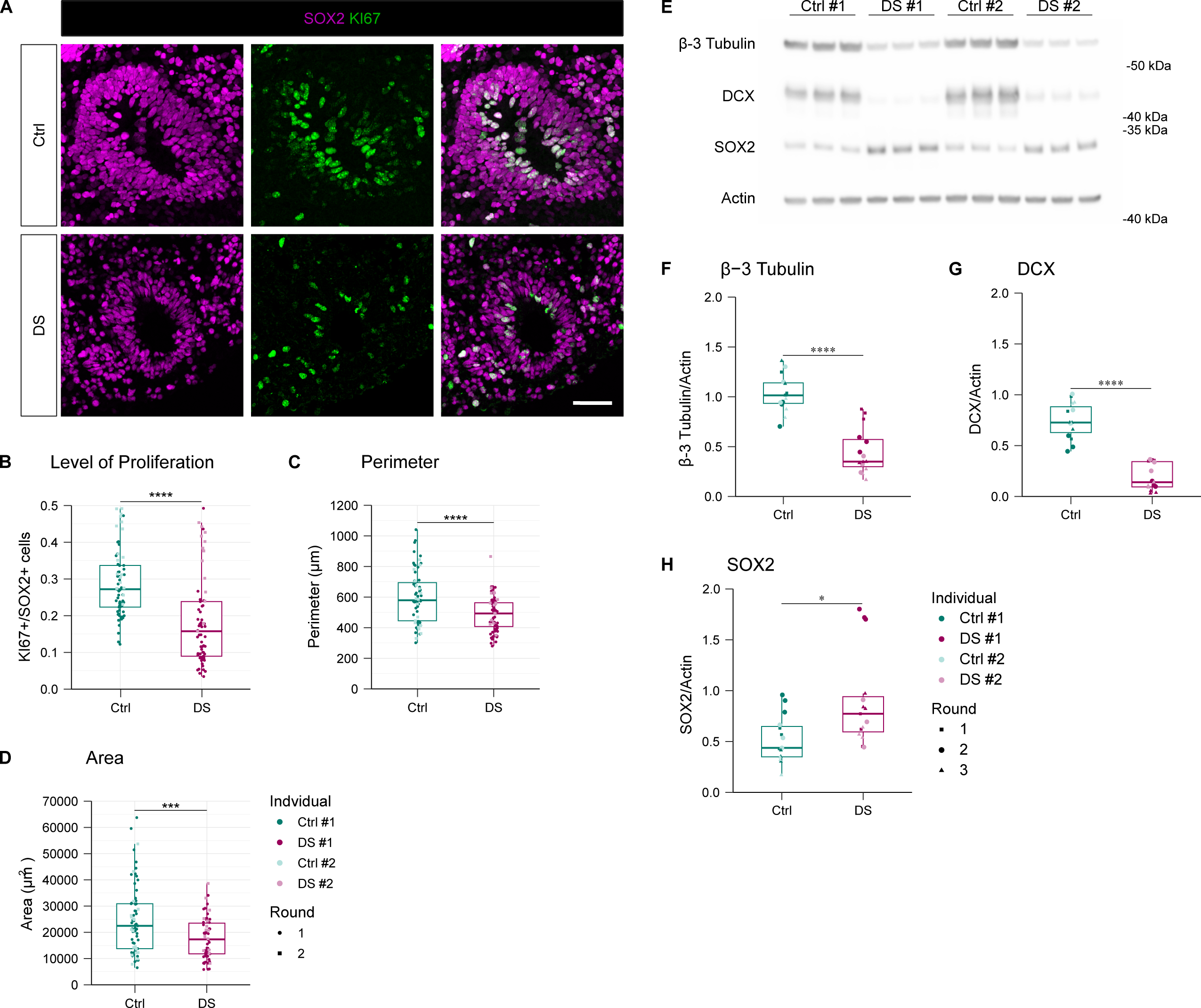
DS proliferative zones exhibit reduced expansion. (A) Representative immunostaining of SOX2 and KI67 from d25 Ctrl and DS dorsal forebrain organoid cryosections. (B-D) Quantification of proliferating cell count (KI67+/SOX2+), neural stem cell count (SOX2+), area (μm^2^), and perimeter (μm) of proliferative zones (pz). (DS/Ctrl (n = 70, 1pz per n, 10-13 orgs, 1 round). Data represented as mean ± SEM. Data analyzed by one-way ANOVA followed by Tukey’s multiple comparisons test. * p ≤ 0.05, ** p ≤ 0.01, *** p ≤ 0.001, **** p ≤ 0.0001. Scale bar, 50 µm. (E) Representative immunoblots of β-3 tubulin, DCX, and SRY SOX2 from d25 Ctrl and DS dorsal forebrain organoids. (F-H) Quantification of protein expression normalized to actin. DS/Ctrl (n = 15, 10 orgs per n, 3 rounds). Data represented as mean ± SEM. Data analyzed by unpaired two-tailed Student’s t-test. * p ≤ 0.05, ** p ≤ 0.01, *** p ≤ 0.001, **** p ≤ 0.0001.

**Figure 3.**
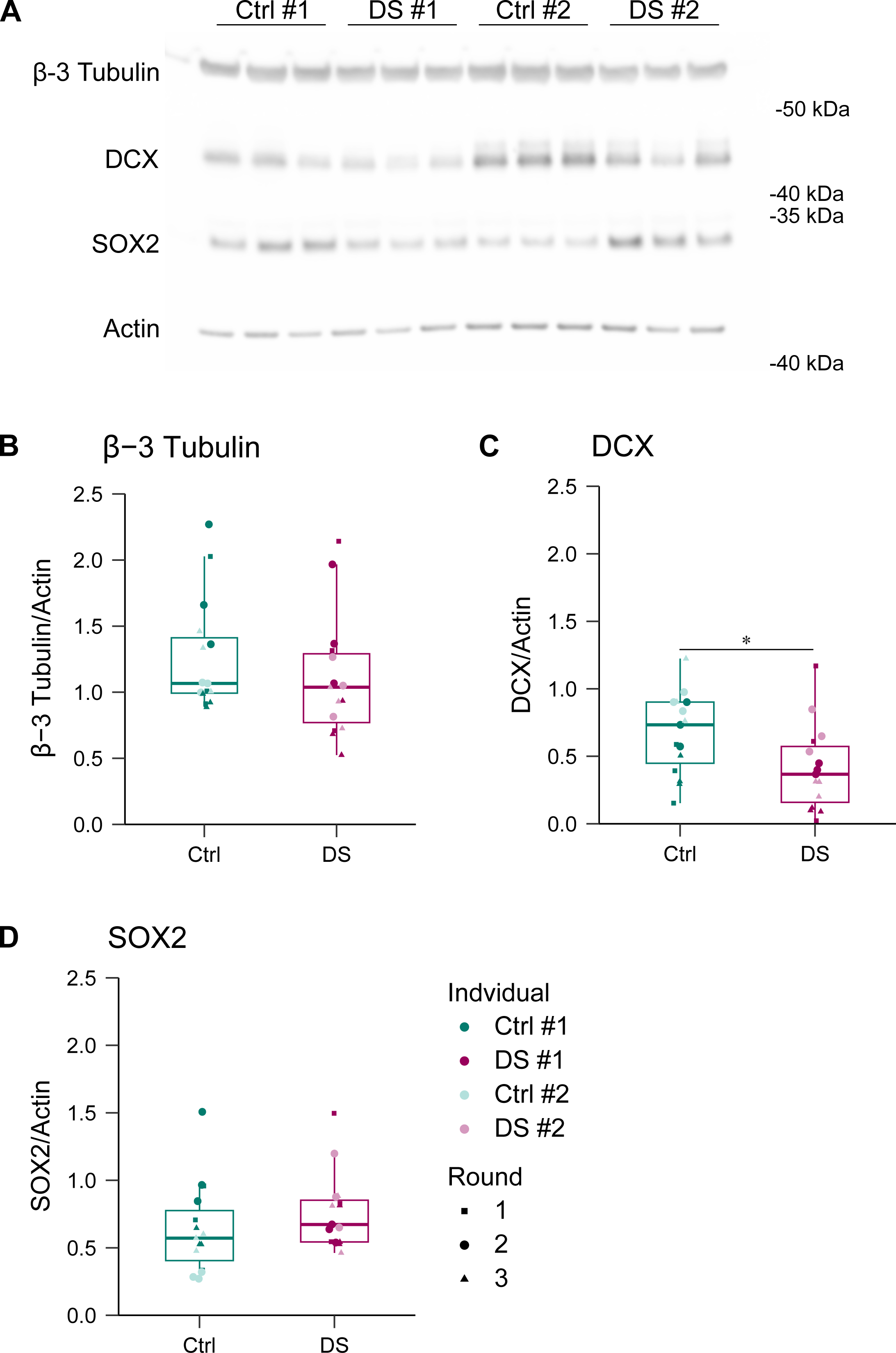
d50 DS cortical organoids exhibit reduced neuroblast protein expression. (A) Representative immunoblots of β-3 tubulin, DCX, and SOX2 from d50 Ctrl and DS dorsal forebrain organoids. (B-D) Quantification of protein expression normalized to actin. DS/Ctrl (n = 15, 10 orgs per n, 3 rounds). Data represented as mean ± SEM. Data analyzed by unpaired two-tailed Student’s t-test. * p ≤ 0.05, ** p ≤ 0.01, *** p ≤ 0.001, **** p ≤ 0.0001.

### Normalization of APP gene copy number reduced APP-FL protein expression and processing in DS cortical organoids

To address the role of increased APP gene dosage in DS corticogenesis, we obtained a genetically modified DS #2 line in which CRISPR-Cas9 was utilized to introduce a stop codon prior to the final exon of a single APP allele with the intent to induce nonsense mediated decay (DS #2 APP +/+/-). A corresponding DS #2 “mock CRISPR-Cas9” (DS#2 MC) line was developed in parallel as a control. Next, we asked if normalization of APP gene copy number is sufficient to reduce APP-FL protein expression in DS cortical organoids. DS #2 APP +/+/- organoids exhibited decreased APP-FL protein levels across all examined time points relative to DS #2 MC (**Figure 4A-D**). In continuity, DS #2 APP +/+/- organoid conditioned media exhibited a significant decrease in Aβ-40 and Aβ-42 levels at d50 and d75 relative to DS #2 MC (**Figure 4E-F**). No alterations in Aβ-42 and Aβ-40 ratio were observed across genotypes (**Figure 4G**).

**Figure 4.**
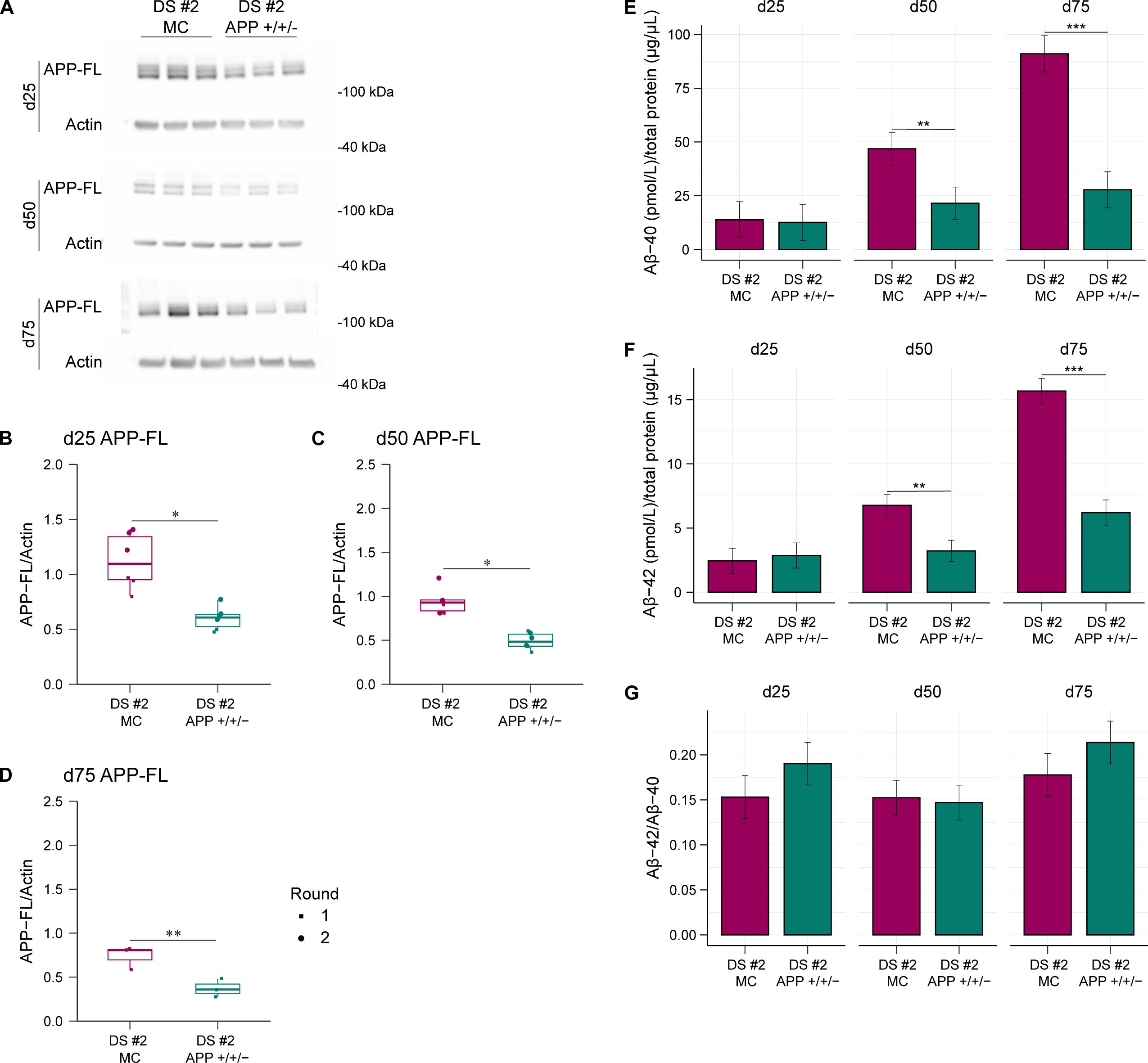
Normalization of APP gene copy number reduced APP-FL protein expression and processing in DS cortical organoids. (A) Representative immunoblots of APP-FL from d25, d50, and d75 DS #2 MC and DS #2 APP +/+/- dorsal forebrain organoids. (B-D) Quantification of protein expression normalized to β-actin. DS #2 MC/APP +/+/- (n= 3-6, 10 orgs per n, 1-2 rounds). Data represented as mean ± SEM. Data analyzed by unpaired two-tailed Student’s t-test. * p ≤ 0.05, ** p ≤ 0.01, *** p ≤ 0.001, **** p ≤ 0.0001. (E-G) Levels of secreted Aβ-40 and Aβ-42 and Aβ-42/Aβ-40 ratio from d25, d50, and d75 DS #2 MC and DS #2 APP +/+/- dorsal forebrain organoid conditioned media measured by ELISA. A linear mixed model (LMM) was used to test for differences in Aβ-40, Aβ-42, or Aβ-42/Aβ-40 ratio. Specifically, day (d25, d50, and d75) and line (DS #2 MC and DS #2 APP +/+/-) were fixed effects and individual and round of differentiation were random effects. Post hoc comparisons were performed comparing levels of differences between line within each time point. Multiple comparisons were corrected for using Tukey’s adjustment. DS #2 MC/APP +/+/- (n= 3-6, 5-10 orgs per n, 1-2 rounds). Data represented as mean ± SEM. * p ≤ 0.05, ** p ≤ 0.01, *** p ≤ 0.001, **** p ≤ 0.0001.

### Reduced expansion and neurogenic protein expression mediated by APP gene overdose in DS cortical organoids

To determine the role of increased APP gene dosage in impaired RGC proliferation and organization, immunostaining was performed on DS #2 MC and DS #2 APP +/+/- organoid cryosections for SOX2 and KI67. DS #2 APP +/+/- proliferative zones exhibited significantly increased proliferating SOX2+ RGCs relative to DS #2 MC (**Figure 5A-B**). In continuity, DS #2 APP +/+/- proliferative zones exhibited increased area and perimeter relative to DS #2 MC (**Figure 5C-D**). To begin interrogating the role of increased APP gene dosage in impaired neurogenesis, we ran immunoblots for the stemness protein SOX2 and neurogenic proteins DCX and β-3 tubulin. At d25, DS #2 APP +/+/- cortical organoids exhibited decreased SOX2 and increased DCX and β-3 tubulin protein expression relative to DS #2 MC (**Figure 5E-H**). To determine if this rescue persisted, we performed the same immunoblotting at d50. DS #2 APP +/+/- cortical organoids exhibited no alterations in SOX2, DCX, and β-3 tubulin protein expression relative to DS #2 MC (**Figure 6A-D**).

**Figure 5.**
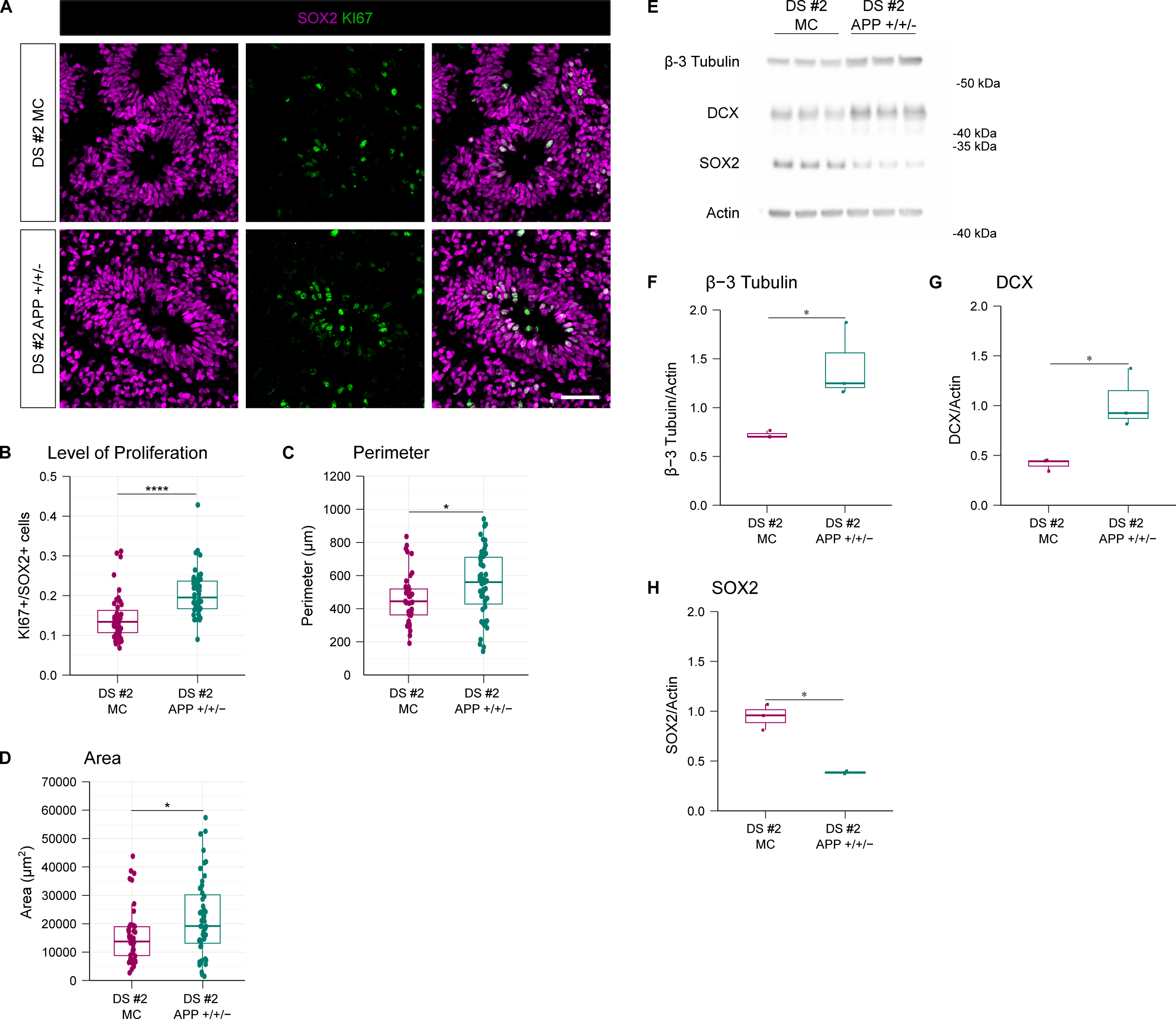
Normalization of APP gene copy number increased proliferative zone expansion in DS cortical organoids. (A) Representative immunostaining of SOX2 and KI67 from d25 DS #2 MC and DS #2 APP +/+/- dorsal forebrain organoid cryosections. (B-D) Quantification of proliferating cell count (KI67+/SOX2+), neural stem cell count (SOX2+), area (μm^2^), and perimeter (μm) of proliferative zones (pz). DS #2 MC/APP +/+/- (n= 50, 1 pz per n, 10 orgs, 1 round). Data represented as mean ± SEM. Data analyzed by one-way ANOVA followed by Tukey’s multiple comparisons test. * p ≤ 0.05, ** p ≤ 0.01, *** p ≤ 0.001, **** p ≤ 0.0001. Scale bar, 50 µm. (E) Representative immunoblots of β-3 tubulin, DCX, and SRY SOX2 from d25 DS #2 MC and DS #2 APP +/+/- dorsal forebrain organoids. (F-H) Quantification of protein expression normalized to actin. DS #2 MC/APP +/+/- (n= 3, 10 orgs per n, 1 round). Data represented as mean ± SEM. Data analyzed by unpaired two-tailed Student’s t-test. * p ≤ 0.05, ** p ≤ 0.01, *** p ≤ 0.001, **** p ≤ 0.0001.

**Figure 6.**
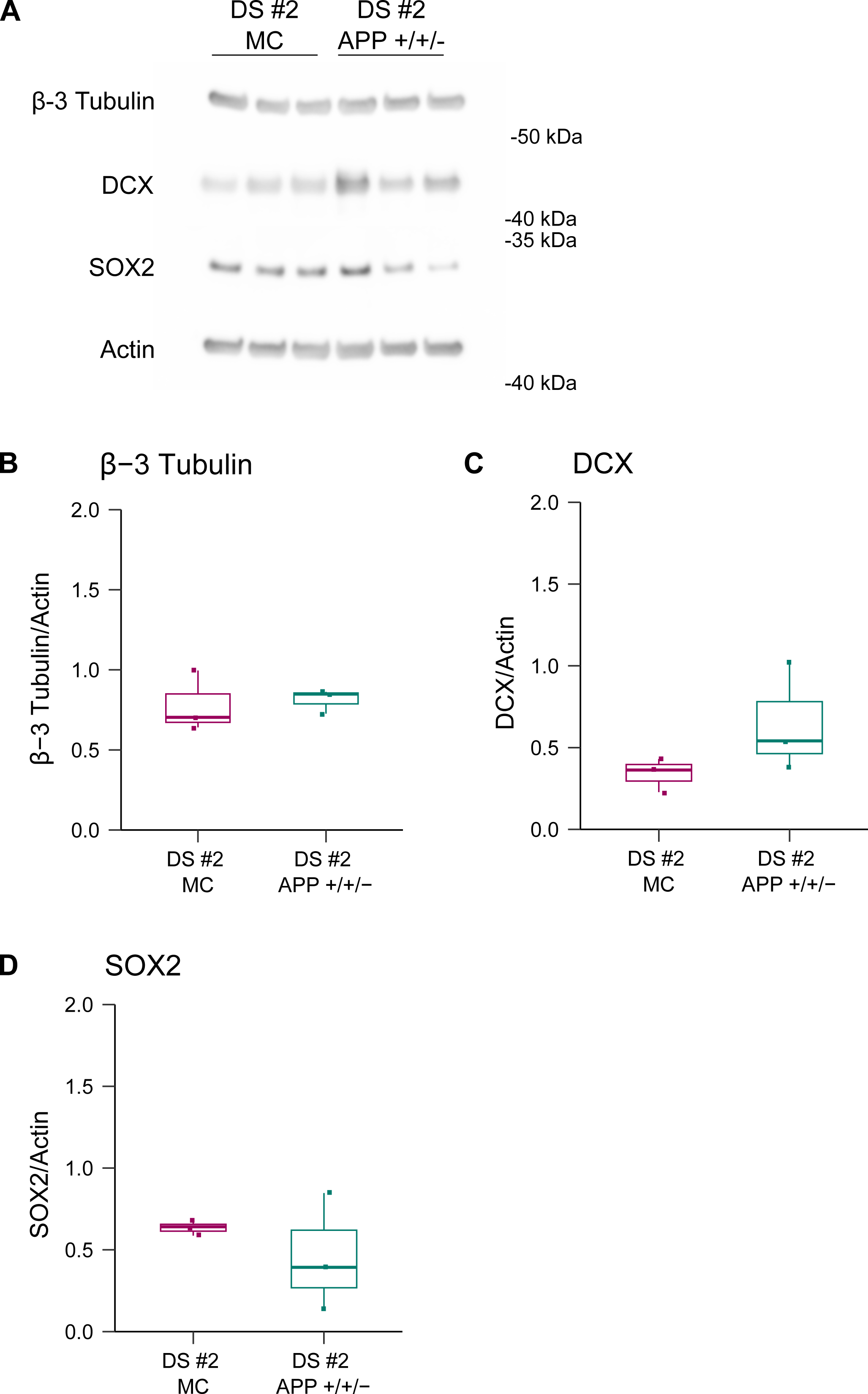
Normalization of APP gene copy number does not alter stem or neurogenic protein expression in d50 DS cortical organoids. (A) Representative immunoblots of β-3 tubulin, DCX, and SOX2 from d50 DS #2 MC and DS #2 APP +/+/- dorsal forebrain organoids. (B-D) Quantification of protein expression normalized to actin. DS #2 MC/APP +/+/- (n= 3, 10 orgs per n, 1 round). Data represented as mean ± SEM. Data analyzed by unpaired two-tailed Student’s t-test. * p ≤ 0.05, ** p ≤ 0.01, *** p ≤ 0.001, **** p ≤ 0.0001.

### Altered transcription mediated by APP gene overdose in DS cortical organoids

Overdose of APP gene copy number has been shown to contribute to transcriptional changes in 2D DS cortical neuron cultures89,114. However, its impact on cell populations of early neurogenesis in DS remains to be investigated. To that end, we performed single-cell RNA sequencing on d25 and d50 cortical organoids across genotypes. Significantly differentially expressed genes (DEGs) with the highest average log_2_ fold change when comparing DS #1/Ctrl #1 and DS #2/Ctrl #2 were identified at d25 in RGCs, intermediate progenitor cells (IPCs), and excitatory neurons (ENs) (**Figure 7A-C**). The same analysis was performed at d50 (**Figure 8A-C**). Significant DEGs with the highest average log_2_ fold change were identified when comparing DS #2 APP +/+/-/DS #2 MC at d25 in RGCs, IPCs, and ENs (**Figure 9A-C**). The same analysis was performed at d50 (**Figure 10A-C**). Next, we performed Gene Ontology (GO) pathway analysis and cluster mapping of significant DEGs. At d25, DS cortical organoids exhibited significantly altered pathways related to RNA processing and cellular respiration (**Figure 7D-F**). These alterations persisted out to d50 (**Figure 8D-F**). At d25, DS #2 APP +/+/- cortical organoids exhibited significantly altered pathways related to RNA processing and autophagy (**Figure 9D-F**). At d50, DS #2 APP +/+/- cortical organoids exhibited significantly altered pathways related to RNA processing, ER stress, and oxidative phosphorylation (**Figure 10D-F**).

**Figure 7.**
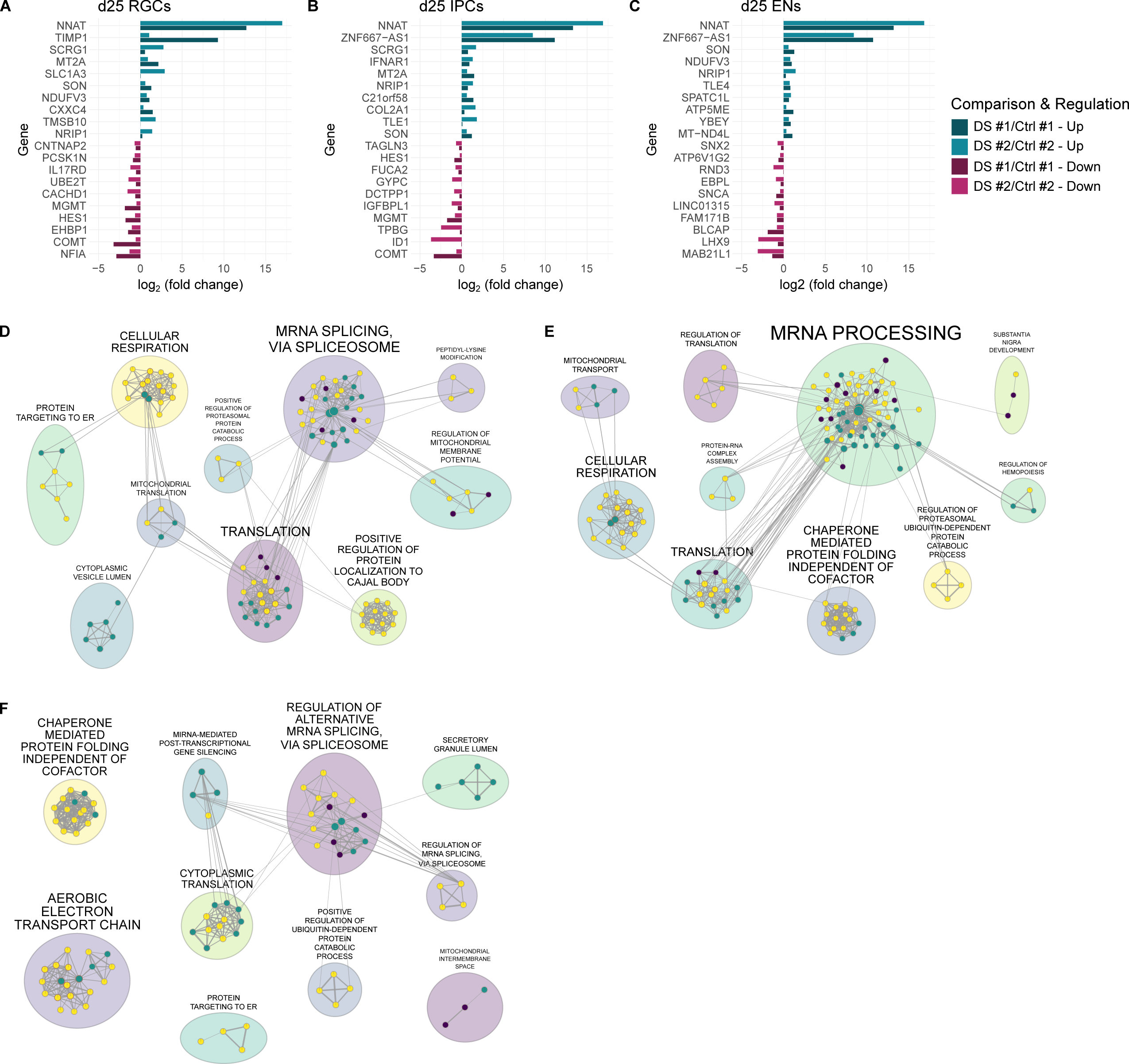
d25 DS cortical organoids exhibit altered RNA processing and cellular respiration pathways. (A-C) Bar plot showing DEGs with a significant cut off value of p < 0.05 and highest average log_2_ fold change when comparing d25 DS#1/Ctrl #1 and DS#2/Ctrl #2 cortical organoids. (D-F) Cytoscape cluster mapping of altered GO pathways for d25 DS and Ctrl cortical organoids. Analysis based on log_2_ fold change ratio of DEGs with a significant cut off value of p < 0.05.

**Figure 8.**
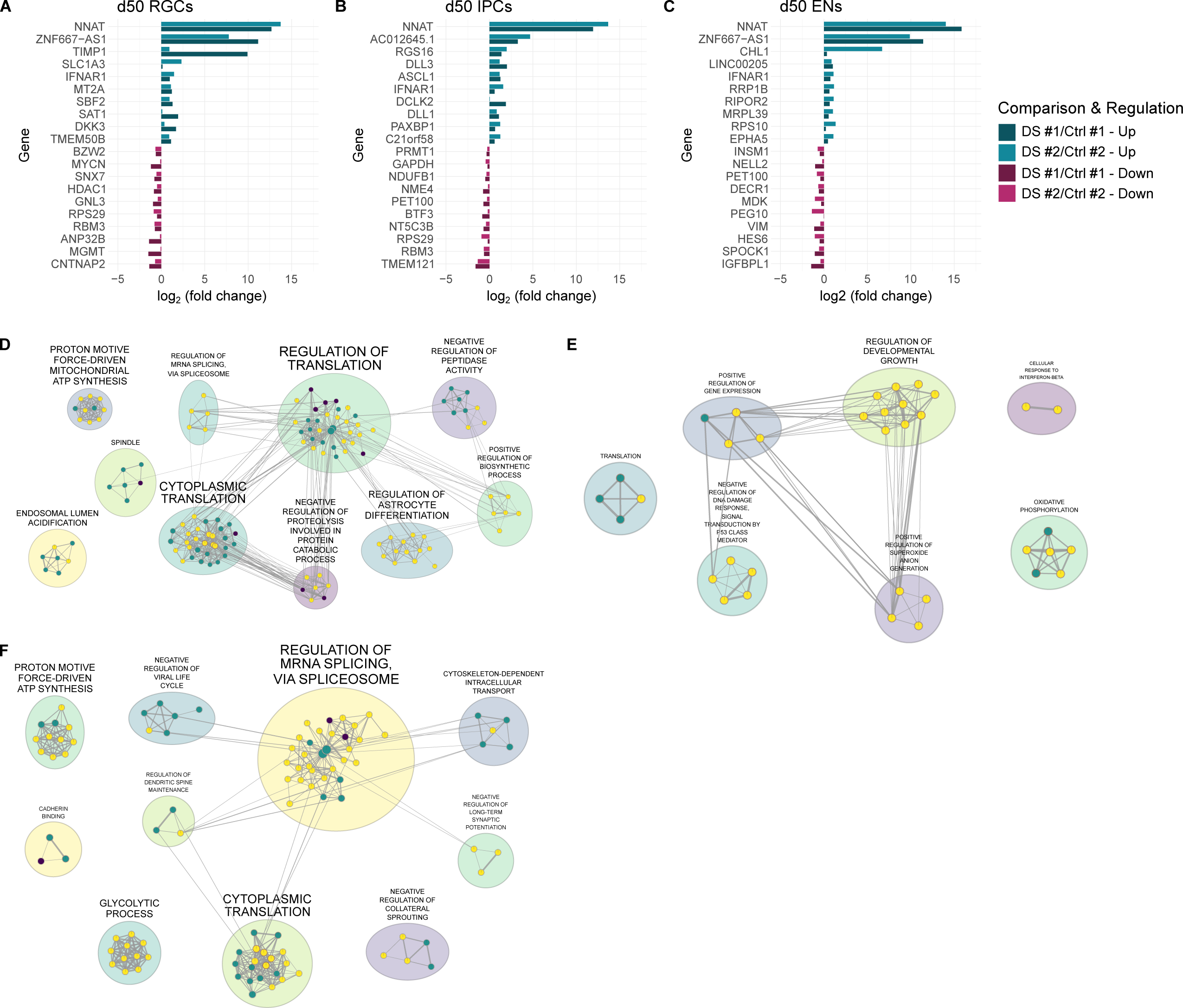
d50 DS cortical organoids exhibit altered RNA processing and cellular respiration pathways. (A-C) Bar plot showing DEGs with a significant cut off value of p < 0.05 and highest average log_2_ fold change when comparing d50 DS#1/Ctrl #1 and DS#2/Ctrl #2 cortical organoids. (D-F) Cytoscape cluster mapping of altered GO pathways for d50 DS and Ctrl cortical organoids. Analysis based on log_2_ fold change ratio of DEGs with a significant cut off value of p < 0.05.

**Figure 9.**
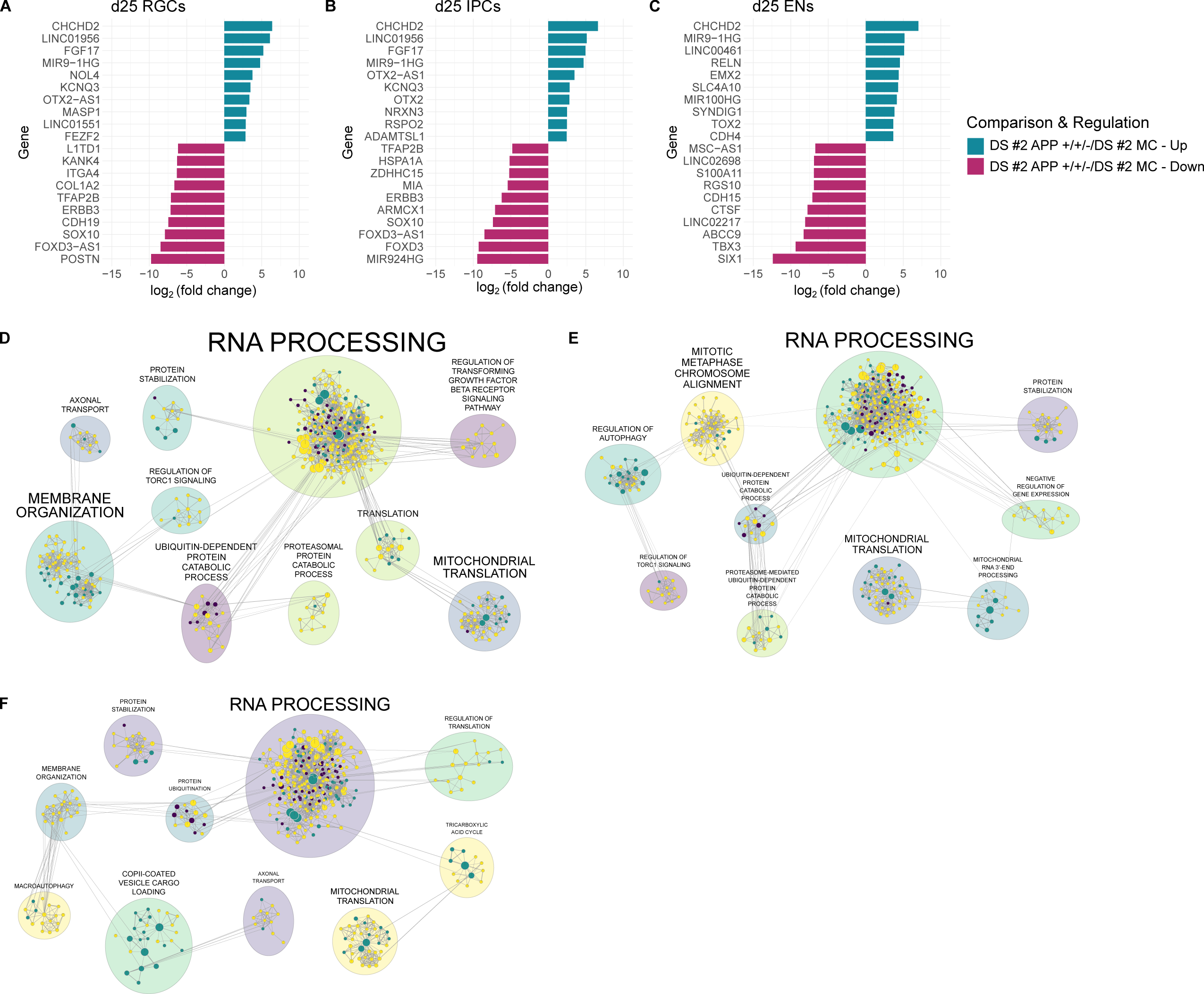
Normalization of APP gene copy number altered RNA processing and autophagy pathways in d25 DS cortical organoids. (A-C) Bar plot showing DEGs with a significant cut off value of p < 0.05 and highest average log_2_ fold change when comparing d25 DS #2 MC and DS #2 APP +/+/- cortical organoids. (D-F) Cytoscape cluster mapping of altered GO pathways for d25 DS #2 MC and DS #2 APP +/+/- cortical organoids. Analysis based on log_2_ fold change ratio of DEGs with a significant cut off value of p < 0.05.

**Figure 10.**
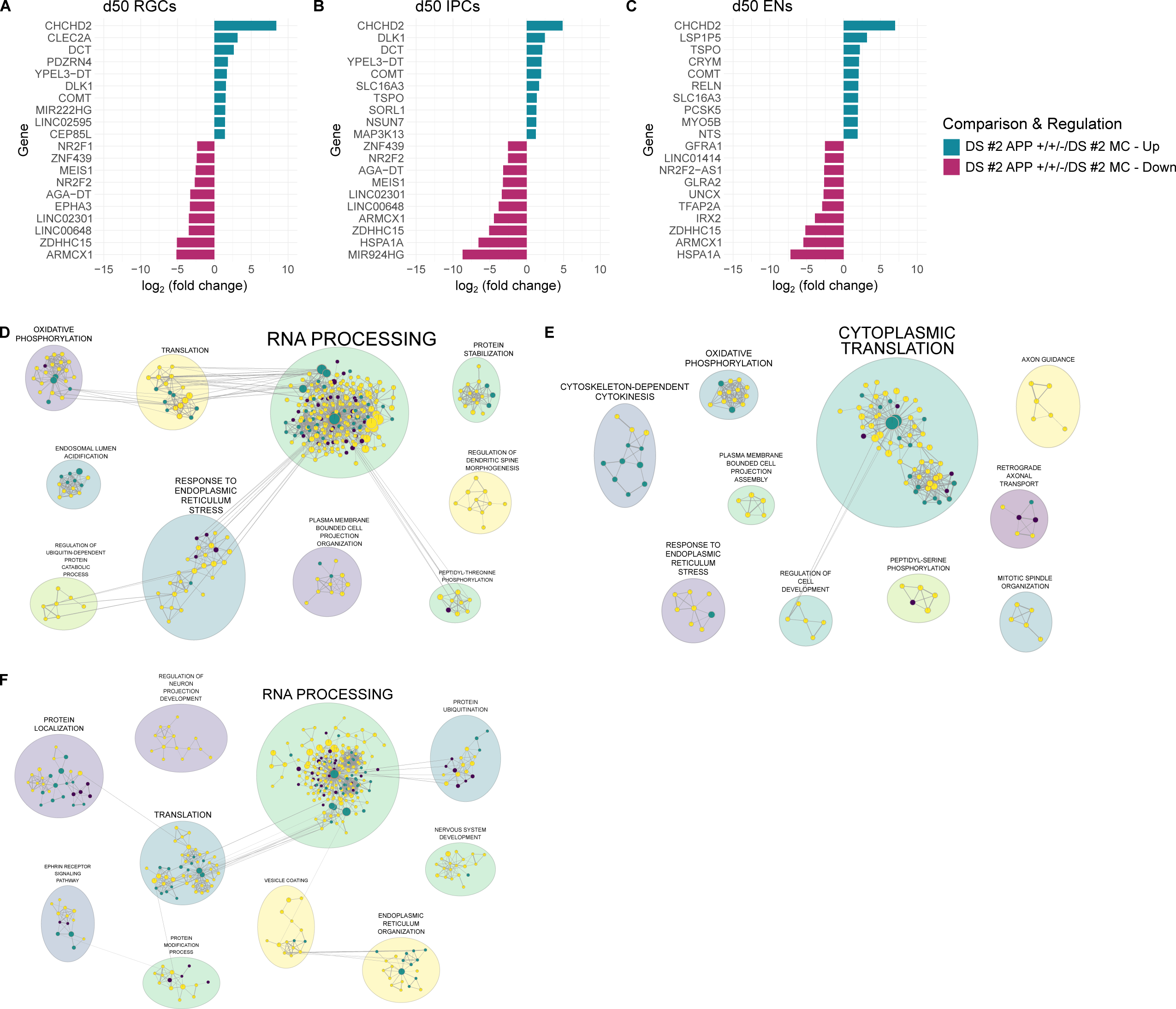
Normalization of APP gene copy number altered RNA processing and autophagy pathways in d50 DS cortical organoids. (A-C) Bar plot showing DEGs with a significant cut off value of p < 0.05 and highest average log_2_ fold change when comparing d50 DS #2 MC and DS #2 APP +/+/- cortical organoids. (D-F) Cytoscape cluster mapping of altered GO pathways for d50 DS #2 MC and DS #2 APP +/+/- cortical organoids. Analysis based on log_2_ fold change ratio of DEGs with a significant cut off value of p < 0.05.

## Discussion

In this study, we found that increased APP-FL protein expression in DS cortical organoids relative to Ctrl coincided with the appearance of IPCs and neurons. This is consistent with previous studies showing that *APP* mRNA expression increased as neurogenesis progressed peaking at neuronal differentiation and neurite outgrowth^46^. Interestingly, limited analysis of DS fetal tissue has shown only an increase in APP mRNA but not APP-FL protein expression ^47–50^. It is of note that protein analysis was done utilizing only immunoblotting of 17 to 19 gestational week DS fetal brains. However, cultured DS NPCs have exhibited significantly increased APP-FL protein expression^51,52^. Given the numerous mechanisms through with APP regulates neurogenesis, rigorous studies delineating which cellular subtypes overexpress APP at what gestational week in DS fetal tissue are needed^31,46,51,53,54^. Our data show that silencing of one APP allele utilizing CRISPR-Cas9 in a DS iPSC line is sufficient to reduce APP-FL protein expression in derived cortical organoids at all examined time points. These results are consistent with previous studies in DS iPSC derived neuronal cultures^35,36^. Notably, silencing of one allele was sufficient to reduce APP-FL protein expression in d25 DS cortical organoids that exhibited similar levels to Ctrl. Given that APP mRNA is overexpressed in DS NPCs, it is likely post-transcriptional or post-translational mechanisms are regulating APP-FL protein expression in early neurogenesis^55,56^. Further investigation of how microRNAs (miRNAs) regulate APP expression in DS may provide insight to the discordant mRNA and protein levels at this time point ^57–59^.

HSA21 triplication results in an early, pervasive accumulation of amyloid pathology. Diffuse Aβ-42 deposits have been reported as early as 8 up to 27 years old, an earlier deposition than in AD^60–63^. Increased soluble Aβ precedes this plaque deposition^64–66^. Higher levels of soluble Aβ-42 were observed in one study at 21 gestational weeks^67^. Rigorous characterization of accumulation of amyloid pathology is necessary for appropriately timed therapeutic interventions. We found that DS cortical organoids as young as d50 secrete increased levels of Aβ-40 and Aβ-42 relative to Ctrl. These results are in line with previously published organoid experiments from additional DS individuals^68,69^. Therefore, increased soluble Aβ may occur as early as mid-gestation in DS. Importantly, we show silencing of one APP allele in DS cortical organoids resulted in a reduction in secreted Aβ. Thus, though multiple HSA21 genes have been shown to modulate amyloid pathology, APP expression likely plays a fundamental role in early increased Aβ secretion^70^.

To investigate NPC pool formation, we characterized proliferative zone cell composition and morphology at d25. We showed that d25 DS proliferative zones contain fewer proliferating RGCs and overall decreased perimeter and area relative to Ctrl. Our results agree with a previously published characterization of DS organoids derived from additional individuals^27^. Limited studies reported decreased proliferation of cells populating both the VZ and SVZ and count of oSVZ SOX2+ RGCs in mid-gestation DS fetal tissue^2,3^. The VZ and SVZ of Ts65Dn mice have exhibited decreased thickness, reduced cell proliferation, and prolonged cell cycle^8^. Thus, reduced proliferation of VZ and SVZ RGCs is a hallmark DS phenotype. We found that silencing one APP allele restored expansion of proliferative zones in d25 DS cortical organoids. Next, we profiled the progression of neurogenesis in cortical organoids. We demonstrated that d25 DS cortical organoids expressed increased stem and decreased neurogenic proteins relative to Ctrl. The decrease in the neuroblast protein DCX persisted out to d50. Increased expression of SOX2, implicated in maintaining self-renewing RGCs, in concert with decreased expression of DCX and β-3 tubulin, implicated in IPC migration and differentiation, suggests deficits in formation and maintenance of the IPC pool in DS cortical organoids^43,44,71–74^. Several groups have shown the DS NPCs suffer from oxidative stress-induced senescence that significantly impacts proliferation and differentiation^5,13–16^. Silencing of one APP allele in d25 DS cortical organoids was sufficient to restore levels of these stem and neurogenic proteins. Notably, we show that normalization of APP gene copy number alters RNA processing, autophagy, ER stress, and oxidative phosphorylation pathways in RGCs and IPCs. Taken together, these results suggest that reduction in APP expression ameliorated NPC deficits in DS cortical organoids.

In conclusion, we show that APP gene overdose mediates deficits in corticogenesis in DS cortical organoids. Future experiments should aim to determine the mechanisms by which APP mediates these deficits.

## Acknowledgments

We thank Dr. Peter Toth and Dr. Ke Ma at the UIC Fluorescence Imaging Core for guidance on microscopy imaging. We thank Dr. Jalees Rehman for guidance on analysis of sequencing data. We thank Dr. Mark Maienschein-Cline and Pinal Kanabar at the UIC Research Informatics Core for analysis of sequencing data. We thank Dr. Zachery Morrissey for guidance on data analysis. This work was financially supported by National Institutes of Health (NIH), National Institute on Aging (NIA) AG079002, AG057468, AG076940, and AG033570 (OL).

## Author Contributions

KR performed the experiments, analyzed the data and wrote the manuscript. DP and LAC assisted in performing experiments. OL supervised the project, revised the manuscript, and provided funding sources (NIA AG079002, AG057468, AG076940, and AG033570).

## Declarations of Interests

The authors declare no conflict of interests.

## Methods

### iPSC Culture

Lines Ctrl #1 (UWWC1-DS2U) and DS #1 (UWWC1-DS1) were obtained from WiCell. Lines Ctrl #2 (ND50027) and DS #2 (ND50026) were obtained from National Institute of Neurological Disorders and Stroke (NINDS)^40,41^. Lines DS #2 MC and DS #2 APP +/+/- were donated by the Young-Pearse lab and have been previously described^75^. iPSCs were cultured on Matrigel hESC-Qualified Matrix (Corning, 354277) in mTeSR Plus (STEMCELL Technologies, 100-0276). Passaging was performed utilizing ReLeSR (STEMCELL Technologies, 100-0483).

### Organoid Culture

Organoids were generated from iPSCs utilizing the STEMdiff Dorsal Forebrain Organoid Differentiation Kit (STEMCELL Technologies, 08620) and STEMdiff Neural Organoid Maintenance Kit (STEMCELL Technologies, 100-0120). Briefly, iPSC colonies were dissociated into single cells utilizing Gentle Cell Dissociation Reagent (STEMCELL Technologies, 100-0485) and 3 million cells were seeded per well into AggreWell 800 24-well plates (STEMCELL Technologies, 34850). Organoids were subsequently maintained according to kit instructions.

### Western Blotting

Organoids were rinsed 3 times for 10 minutes with Dulbecco’s phosphate-buffered saline (DPBS) with no calcium or magnesium (Thermo Fisher Scientific, 14190144) to remove excess media. DPBS was aspirated completely, and organoids were snap frozen in liquid nitrogen. 10 day 25 or 5 day 50 or 75 organoids were pooled together per sample. Organoids were lysed in radio-immunoprecipitation assay (RIPA) buffer (Boston Bio Products, BP-115) with protease and phosphatase inhibitor cocktail (Thermo Fisher Scientific, 78440) by running 10 times through a syringe (Fisher Scientific, BD 305620) and sonicating on ice at 20% power 3 times for 15 s with 5 s rest between. Lysate was centrifuge at 16 thousand g for 15 minutes at 4°C, and the supernatant was transferred to a fresh tube. The concentration of the protein lysate was determined utilizing the Pierce BCA Protein Assay Kit (Thermo Fisher Scientific, 23235). Lysates were prepared in Bolt LDS Sample Buffer (Thermo Fisher Scientific, B0007) and Bolt Sample Reducing Agent (Thermo Fisher Scientific, B0009). Western blot samples were boiled for 5 minutes at 95°C and separated on Bolt Bis-Tris Plus polyacrylamide gels (Thermo Fisher Scientific, NW04125BOX) in Bolt MES SDS Running Buffer (Thermo Fisher Scientific, B0002). The iBlot 2 Dry Blotting System (Invitrogen, IB21001) was utilized to transfer protein from gels to iBlot 2 Transfer Stacks (Thermo Fisher Scientific, IB23002). Nitrocellulose membranes were blocked for 1 hour in 5% non-fat dry milk (milk) in tris-buffered saline containing 0.1% Tween-20 (TBST). Blots were incubated overnight at 4°C in primary antibodies diluted in 5% milk in TBST and subsequently washed 3 times for 15 minutes in TBST. Blots were then incubated for 2 hours in secondary antibodies diluted in 5% milk in TBST and subsequently washed 3 times for 15 minutes in TBST. Membranes were developed utilizing SuperSignal West Pico PLUS Chemiluminescent Substrate (Thermo Fisher Scientific, 34577) and imaged on Azure Biosystems 300q Image Western Blot Imaging System (Azure Biosystems, AZI300-01). Fiji (NIH) was utilized to quantify band intensities. Antibodies utilized in experiments are listed in Appendix A.

### Immunocytochemistry

Organoids were rinsed 3 times for 10 minutes in DPBS to remove excess media and incubated overnight in 4% PFA at 4°C. They were then rinsed 3 times for 10 minutes in DPBS and incubated with 30% sucrose for approximately 24 to 48 hours at 4°C until they sunk to the bottom of the Eppendorf tube. Organoids were incubated for 1 hour at 37°C and snap frozen in a mold utilizing an isopentane bath chilled with liquid nitrogen in a solution of 10% sucrose and 7.5% gelatin in DPBS. Molds were cryosectioned into 20 μm sections and mounted on Superfrost Plus Microscope Slides (Fisher Scientific, 12-550-15). Sections were washed with phosphate-buffered saline containing 0.1% Tween-20 (PBST) for 30 minutes at 37°C to remove excess gelatin and blocked in 0.3% Triton X-100, 2% bovine serum albumin (BSA), and 5% normal donkey serum (NDS) diluted in PBS for 1 hour. After blocking, they were incubated in primary antibodies diluted in blocking buffer overnight at 4°C in a humidity chamber. Sections were washed 3 times for 5 minutes each in PBST and incubated in secondary antibodies diluted in PBST for 2 hours. Sections were then washed 3 times for 5 minutes each in PBST and coverslips were mounted onto slides with ProLong Gold Antifade Mountant with DAPI (Thermo Fisher Scientific, P36931). Antibodies utilized in experiments are listed in Appendix A. Images were acquired at 40x magnification utilizing confocal microscopy (Zeiss LSM 710).

### ELISA

Organoids were rinsed 3 times in DPBS and placed in 1 mL of fresh media. 10 organoids for day 25 or 5 organoids for day 50 or 75 were pooled per well. Organoids were incubated in media for 48 hours. Resulting organoid conditioned media was centrifuged at 2 thousand rpm for 10 minutes at 4°C and supernatant was snap frozen in liquid nitrogen. Protein was extracted from organoids in corresponding wells and total protein concentration was determined as described in Section 2.2.3. To measure Aβ isoforms in organoid conditioned media, Human β Amyloid (1-40) ELISA Kit Wako II (Fujifilm Irving, 298-64601) and Human β Amyloid(1-42) ELISA Kit Wako (Fujifilm Irving, 298-62401) were utilized. Samples were prepared according to kit instructions at a dilution of 1:3 for Aβ-40 and 1:2 for Aβ-42. Resulting concentrations were normalized to total protein extracted from organoids in corresponding well. To measure tau isoforms in organoid protein lysate, PathScan RP Phospho-Tau (Ser396) Sandwich ELISA Kit (Cell Signaling, 43932) and FastScan Total Tau ELISA Kit (Cell Signaling, 57519) were utilized. All lysates were prepared according to kit instructions at a standard protein amount.

### Single-cell RNA Sequencing

Cell number and viability were analyzed using Nexcelom Cellometer Auto2000 with AOPI fluorescent staining method. Sixteen thousand cells were loaded into the Chromium iX Controller (10X Genomics, PN-1000328) on a Chromium GEM-X Single cell 3’ Chip Kit v4 (10x Genomics, PN-1000215), and processed to generate single cell gel beads in the emulsion (GEM) according to the manufacturer’s protocol. The cDNA and library were generated using the GEM-X Single cell 3’ Kit v4 (10x Genomics, PN-1000691) and Dual Index Kit TT Set A (10X Genomics, PN-1000215) according to the manufacturer’s manual. Quality control for constructed library was performed by Agilent Bioanalyzer High Sensitivity DNA kit (Agilent Technologies, 5067-4626) and Qubit DNA HS assay kit for qualitative and quantitative analysis respectively. The multiplexed libraries were pooled and sequenced on Illumina Novaseq X Plus sequencer with 100 cycle kits using the following read length: 28 bp Read1 for cell barcode and UMI, and 90 bp Read2 for transcript. The single cell library preparation and sequencing was done at Northwestern University NUseq facility core.

### Preprocessing

Single-cell gene expression was quantified using CellRanger (10X Genomics). Single cells were filtered based on the following criteria: at least 500 genes expressed, at least 2000 UMI counts over all genes, and less than 15% mitochondrial gene expression. Gene expression was normalized using NormalizeExpression in Seurat^76^. To identify cell types, we used the Seurat label transfer functions FindTransferAnchors and TransferData with 30 principal components, using single-cell data published by Kanton *et al.* as the reference data set with the following modifications: astrocytes and microglia were removed from the reference prior to label transfer, and after label transfer all cells with “Glyc” (glycolysis) labels were removed from our data set. Integrated UMAP coordinates were computed using the IntegrateLayers function from Seurat with the CCAIntegration method^76,77^.

### Differential Expression Analysis

Differential expression statistics between treatment groups within each cell line were computed using FindMarkers from Seurat with the Wilcox test^76^. DS vs Ctrl comparisons were run separately for our two cell lines (Ctrl #1/DS #2 and Ctrl #2/DS #2), and we then performed a correspondence analysis to identify concordant results from the two comparisons: a correspondence statistic was computed as the geometric mean of the -log10 adjusted p-values, times the sign of the product of the log2 fold-changes. Adjusted p-values < 1e-10 were set to a floor of 1e-10 first. We then filtered for highly concordant genes based on correspondence > 4, ensuring that we only preserved effects that were statistically significant in both cell lines, and changing in the same direction. To further compare with genes altered in the DS #2 APP +/+/- vs DS #2 MC, we identified genes with opposite effects in DS #2 APP +/+/-/DS #2 MC relative to the DS/Ctrl comparison. Pathway enrichment statistics were computed against Gene Ontology Biological Process (GO BP) pathways obtained from MSigDB for statistically significant or concordance-filtered genes using Fisher’s Exact test in R^76^.

### Statistics

Statistical analysis was performed utilizing R version 4.3.1 (2024-06-16). One-way analysis of variance (ANOVA) and linear mixed model (LMM) analysis were performed utilizing emmeans and lmerTest packages in R respectively. Plots were visualized utilizing ggplot2 package in R and modified utilizing Inkscape version 1.2.1 (2022-07-14). In all graphs, data is shown as mean ± SEM. The following was used for p values: ns > 0.05, *p < 0.05, **p < 0.01, ***p < 0.001, and ****p < 0.0001.

## Notes

### Competing Interest Statement

The authors have declared no competing interest.

